# Diversity of SARS-CoV-2 genome among various strains identified in Lucknow, Uttar Pradesh

**DOI:** 10.1101/2021.10.05.463185

**Authors:** Biswajit Sahoo, Pramod Kumar Maurya, Ratnesh Kumar Tripathi, Jyotsana Agarwal, Swasti Tiwari

## Abstract

Severe acute respiratory syndrome coronavirus 2 (SARS-CoV-2) has emerged as a significant challenge worldwide. Rapid genome sequencing of SARS-CoV-2 is going on across the globe to detect mutations and genomic modifications in SARS-CoV-2. In this study, we have sequenced twenty-three SARS-CoV-2 positive samples collected during the first pandemic from the state of Uttar Pradesh, India. We observed a range of already reported mutations (2-22), including; D614G, L452R, Q613H, Q677H, T1027I in the S gene; S194L in the N gene; Q57H, L106F, T175I in the ORF3. Few unreported mutations such as P309S in the ORF1ab gene; T379I in the N gene; and L52F, V77I in the ORF3a gene were also detected. Phylogenetic genome analysis showed similarity with other SARS-CoV-2 viruses reported from Uttar Pradesh. The observed mutations may be associated with SARS-CoV-2 virus pathogenicity or disease severity.

## Introduction

In December 2019, Wuhan, Hubei province, China first reported numerous pneumonia-like cases with unidentified etiology. Later it was identified as a novel coronavirus (COVID-19)[1]. According to WHO, as of now 13 August 2021, a total of 205,338,159 confirmed COVID-19 cases, including 4,333,094 deaths have been reported worldwide and 32,117,826 Confirmed Cases and 430,254 deaths in India. The state of Uttar Pradesh witnessed 17, 08,876 confirmed cases and 22,782 deaths according to the Government of India. The first strain of Wuhan-Hu-1 coronavirus was isolated and the sequenced complete genome was 29.9 kb[2]. Also, other types of coronavirus including SARS-CoV and MERS-CoV have been identified previously, which infect humans which are positive-sense RNA genomes with 27.9 kb and 30.1 kb, respectively[3]. To understand the genetic variants of SARS-CoV-2, genome sequencing is an essential tool to track cases and determining microbial provenance[4], [5]. The first SARS-CoV-2 genome sequence was publicly available on January 10, 2020 (GenBank ID: MN908947.3)[2]. Since then multiple sequences have been submitted to publicly available databases such as GeneBank and GISAID globally. Study of this extensive genomic sequencing to identify the mutations which can increase transmissibility and virulence of the virus[6], [7]. SARS-CoV-2 is an enveloped positive-sense single-stranded RNA virus that has multiple genes which code for different proteins such as open reading frames, including ORF1a, ORF1b ORF3a, ORF6, ORF7a, ORF7b, ORF8, and ORF10; S gene (Surface glycoprotein), N gene (nucleocapsid phosphoproteins), M gene (membrane glycoprotein) and E gene (envelope protein)[8]. Viral genome of SARS-CoV-2, mainly placed in the first ORF (ORF1a/b) and translated into two replicase polyproteins (pp1a and pp1ab), 16 non-structural proteins (NSP), and RNA-dependent RNA polymerase which is important for replication and survival in the host[9]. The remaining ORFs such as ORF3a, ORF7, and ORF8 genes code for accessory and structural proteins, however, their role is not yet fully understood. ORF3a in SARS-CoV has played a significant role in viral release to the host[10], [11]. The exact role of ORF7 and ORF8 is still unknown but several reports suggested that they are involving in viral replication and have immune responses[12], [13]. Open-source databases such as the GenBank-NCBI and GISAID have a huge number of SARS-CoV-2 genomic sequencing data to identify the mutation, SNPs (single nucleotide polymorphism) in SARS-CoV-2. SNPs in the SARS-CoV-2 genome lead to the missense variant such as P323L, P4715L in ORF1ab, D614G, N439K in S gene, R203K, R202K, and G204R in N gene which is commonly reported. P323L, P4715L mutation in ORF1ab has played an important role in regulating RNA-dependent RNA polymerase[14]. D614G, N439K mutation in S gene has associated with increase infectivity[15]. R203K, R202K, and G204R mutations in the N gene are linked with viral survival and replication in the host[16]. Knowing the variants which could escape the immunity, by genome sequencing is essential to stop the coronavirus across the globe. In this study, we sequenced the SARS-CoV-2 genome from 24 SARS-CoV-2 positive samples received at Dr. Ram Manohar Lohia Institute of Medical Sciences, Lucknow during the first peak during the pandemic in the state of Uttar Pradesh, India. Various mutations were observed in the genomic sequences including previously reported mutations as well as novel mutations.

## Materials and Methods

### Collection of Covid-19 positive RNA Samples

RNA from twenty-three Covid-19 positive samples was obtained from Dr. Ram Manohar Lohia Institute of Medical Sciences, Lucknow. The presence of SARS-CoV-2 was detected by COVID-19 RT-qPCR kit ((Labgun, lab, Genomics. co. Ltd, Republic of Korea). Cq values between 18-35 were taken for sequencing. This study protocol was approved by Institutional Human Ethics Committee SGPGIMS, Lucknow (Ref N. 111 PGI/BE/327/2020)

### Whole-genome sequencing

For sequencing, libraries were constructed using a ligation kit (SQK-LSK109) as described in PCR-tiling of COVID-19 virus protocol (PTC_9096_v109_revF_06Feb2020; Oxford Nanopore Technologies). Briefly, 23 SARS-CoV-2 positive RNA samples were isolated from swabs positive for the presence of SARS-CoV-2 in RT-qPCR assay (quantification cycle (Cq) values18-31;(Table 1) and were converted into complementary DNA (cDNA). Then the cDNA products were amplified using the primer pools spanning the SARS-CoV-2 whole genome sequence (i.e., 400-bp Artic nCoV-2019 V3 panel (https://github.com/artic-network/artic-ncov2019) purchased from Integrated DNA Technologies according to the manufacturer’s instructions. DNA library preparation (SQK-LSK-109, Oxford Nanopore Technologies, United Kingdom), purification using AMPure XP magnetic beads (Beckman Coulter), adaptor ligation, and barcoding EXP-NBD104 (barcodes 1-12) or EXP-NBD114 (barcodes 13-24) kits (Oxford Nanopore Technologies, United Kingdom) were done as per the manufacturer’s instructions. DNA libraries were pooled and loaded on the R9.4.1 flow cell (FLO-MIN106, Oxford Nanopore Technologies, United Kingdom). The sequencing was performed using a MinION Mk-1b device (Oxford Nanopore Technologies).

**Table 1:** Cq values of the isolated samples for SARS-CoV-2 detection.

| Sample ID | Age | Sex | Cq values |  |  |
| --- | --- | --- | --- | --- | --- |
|  |  |  | E gene | N gene | RdRp |
| hCoV-19/India/UP-SGPGI-01/2020 | 39 | M | 23.14 | 24.37 | 23.64 |
| hCoV-19/India/UP-SGPGI-02/2020 | 45 | M | 22.64 | 23.68 | 21.93 |
| hCoV-19/India/UP-SGPGI-03/2020 | 29 | M | 25.46 | 24.48 | 22.48 |
| hCoV-19/India/UP-SGPGI-04/2020 | 39 | M | 26.78 | 26.14 | 26.98 |
| hCoV-19/India/UP-SGPGI-06/2020 | 23 | M | 28.42 | 28.03 | 28.56 |
| hCoV-19/India/UP-SGPGI-07/2020 | 57 | M | 18.93 | 19.43 | 19.13 |
| hCoV-19/India/UP-SGPGI-08/2020 | 48 | M | 29.04 | 29.46 | 29.44 |
| hCoV-19/India/UP-SGPGI-09/2020 | 47 | M | 27.78 | 27.16 | 27.04 |
| hCoV-19/India/UP-SGPGI-10/2020 | 71 | M | 27.56 | 27.86 | 27.13 |
| hCoV-19/India/UP-SGPGI-11/2020 | 72 | M | 28.92 | 28.48 | 27.89 |
| hCoV-19/India/UP-SGPGI-12/2020 | 50 | M | 29.06 | 28.06 | 29.65 |
| hCoV-19/India/UP-SGPGI-13/2020 | 39 | F | 29.56 | 28.18 | 29.03 |
| hCoV-19/India/UP-SGPGI-14/2020 | 40 | F | 31.27 | 30.56 | 29.25 |
| hCoV-19/India/UP-SGPGI-15/2020 | 28 | F | 24.06 | 24.56 | 23.94 |
| hCoV-19/India/UP-SGPGI-16/2020 | 37 | F | 26.84 | 26.12 | 25.68 |
| hCoV-19/India/UP-SGPGI-17/2020 | 19 | F | 24.04 | 23.68 | 23.55 |
| hCoV-19/India/UP-SGPGI-18/2020 | 36 | F | 17.88 | 18.12 | 18.36 |
| hCoV-19/India/UP-SGPGI-19/2020 | 48 | F | 26.67 | 26.44 | 26.21 |
| hCoV-19/India/UP-SGPGI-20/2020 | 62 | F | 21.27 | 21.36 | 26.68 |
| hCoV-19/India/UP-SGPGI-21/2020 | 48 | F | 18.15 | 19.34 | 19.65 |
| hCoV-19/India/UP-SGPGI-22/2020 | 49 | F | 28.79 | 28.41 | 27.86 |
| hCoV-19/India/UP-SGPGI-23/2020 | 58 | F | 30.04 | 29.67 | 29.16 |
| hCoV-19/India/UP-SGPGI-24/2020 | 61 | F | 30.86 | 30.64 | 30.78 |

### Genome Assembly, Alignment, and Phylogenetic Analysis

Nanopore sequencing data were base called and de-multiplexed using Guppy v.3.4.4. Variant analysis was performed using Artic analysis pipeline v.1.1.3. (https://artic.network/ncov-2019/ncov2019-bioinformatics-sop.html) using recommended settings. Minimum and maximum read lengths in the Artic guppyplex filter were set to 400 and 700 for the 400-bp amplicons. Wuhan Hu-1 (GenBank ID: MN908947.3) used as reference genome.

MAFFT was used to align the whole sequences of the 23 genomes to the SARS-CoV-2 reference genome (MN908947.3)[17], [18]. SnpEff v4.3 was used to identify the SNPs and changes in the amino acid produced by the gene in the genome[19].

Phylogenetic analysis was done using Nextstrain and aligned using MAFFT v7.471 to assess previously reported genomes. Maximum likelihood trees were generated and identified clade as well as lineages to classify the identified variants using Nextclade [20].

## Results and Discussion

Twenty-three RT-PCR SARS-CoV-2 positive samples from different periods of the pandemic in Uttar Pradesh were selected for this study. Genome coverage was obtained in between 1000X to 8500X for all the samples at the end of sequencing. Next consensus genomes were obtained after assembling with the reference genome. All the sequenced genomes were submitted to GISAID [Table 2].

**Table 2:** Genome similarity of twenty four sequenced genomes when compared to NCBI reference genome (MN908947.3) along with total reads generated and genome coverage obtained.

| **Sample ID** | **GISAID ID** | **Total of Reads** | **Genome Coverage**<br>**Obtained** | **Genome**<br>**Similarity** |
| --- | --- | --- | --- | --- |
| hCoV-19/India/UP-SGPGI-01/2020 | EPI_ISL_4768146 | 47548 | 3275X | 95.4 |
| hCoV-19/India/UP-SGPGI-02/2020 | EPI_ISL_4769080 | 87770 | 5329.6X | 95.5 |
| hCoV-19/India/UP-SGPGI-03/2020 | EPI_ISL_4769421 | 107414 | 5906.7X | 95.4 |
| hCoV-19/India/UP-SGPGI-04/2020 | EPI_ISL_4769787 | 182878 | 9977.6X | 95.6 |
| hCoV-19/India/UP-SGPGI-06/2020 | EPI_ISL_4770946 | 137188 | 6826.2X | 95.5 |
| hCoV-19/India/UP-SGPGI-07/2020 | EPI_ISL_4771322 | 90296 | 5151.2X | 95.5 |
| hCoV-19/India/UP-SGPGI-08/2020 | EPI_ISL_4771568 | 108639 | 6172.8X | 95.5 |
| hCoV-19/India/UP-SGPGI-09/2020 | EPI_ISL_4771854 | 98729 | 5114.9X | 95.6 |
| hCoV-19/India/UP-SGPGI-10/2020 | EPI_ISL_4772199 | 89936 | 5347.3X | 95.6 |
| hCoV-19/India/UP-SGPGI-11/2020 | EPI_ISL_4772715 | 116880 | 5986.6X | 95.5 |
| hCoV-19/India/UP-SGPGI-12/2020 | EPI_ISL_4773080 | 98995 | 5832.5X | 95.5 |
| hCoV-19/India/UP-SGPGI-13/2020 | EPI_ISL_4773776 | 77121 | 5003.9X | 95.5 |
| hCoV-19/India/UP-SGPGI-14/2020 | EPI_ISL_4775580 | 143542 | 8404.1X | 95.6 |
| hCoV-19/India/UP-SGPGI-15/2020 | EPI_ISL_4777224 | 69930 | 3897.6X | 95.6 |
| hCoV-19/India/UP-SGPGI-16/2020 | EPI_ISL_4778343 | 86119 | 4998.6X | 95.5 |
| hCoV-19/India/UP-SGPGI-17/2020 | EPI_ISL_4778982 | 92708 | 5211.1X | 95.6 |
| hCoV-19/India/UP-SGPGI-18/2020 | EPI_ISL_4779221 | 83562 | 4598.4X | 95.6 |
| hCoV-19/India/UP-SGPGI-19/2020 | EPI_ISL_4779775 | 34542 | 2631.8X | 95.6 |
| hCoV-19/India/UP-SGPGI-20/2020 | EPI_ISL_4780686 | 64034 | 3427.8X | 95.4 |
| hCoV-19/India/UP-SGPGI-21/2020 | EPI_ISL_4781620 | 17817 | 1257.8X | 94.6 |
| hCoV-19/India/UP-SGPGI-22/2020 | EPI_ISL_4782798 | 108963 | 6378.9X | 95.6 |
| hCoV-19/India/UP-SGPGI-23/2020 | EPI_ISL_4783944 | 26242 | 1560.3X | 95.5 |
| hCoV-19/India/UP-SGPGI-24/2020 | EPI_ISL_4784200 | 84284 | 5123.9X | 95.6 |

The twenty-three viral SARS-CoV-2 genome assemblies were aligned to the reference genome (SARS-CoV-2 Wuhan-Hu-1 MN908947.3) using MAFFT and genome similarities to the reference genome were calculated (Table 2). All genomes showed more than 95% similarity. Individual genomes were aligned to the NCBI reference genome to predict the mutations in the genome. Using the SnpEff v4.3 tool, various synonymous and missense mutations were detected. A range of 8 to 22 mutations was detected in major genes in twenty-three samples (Table 3).

**Table 3:**
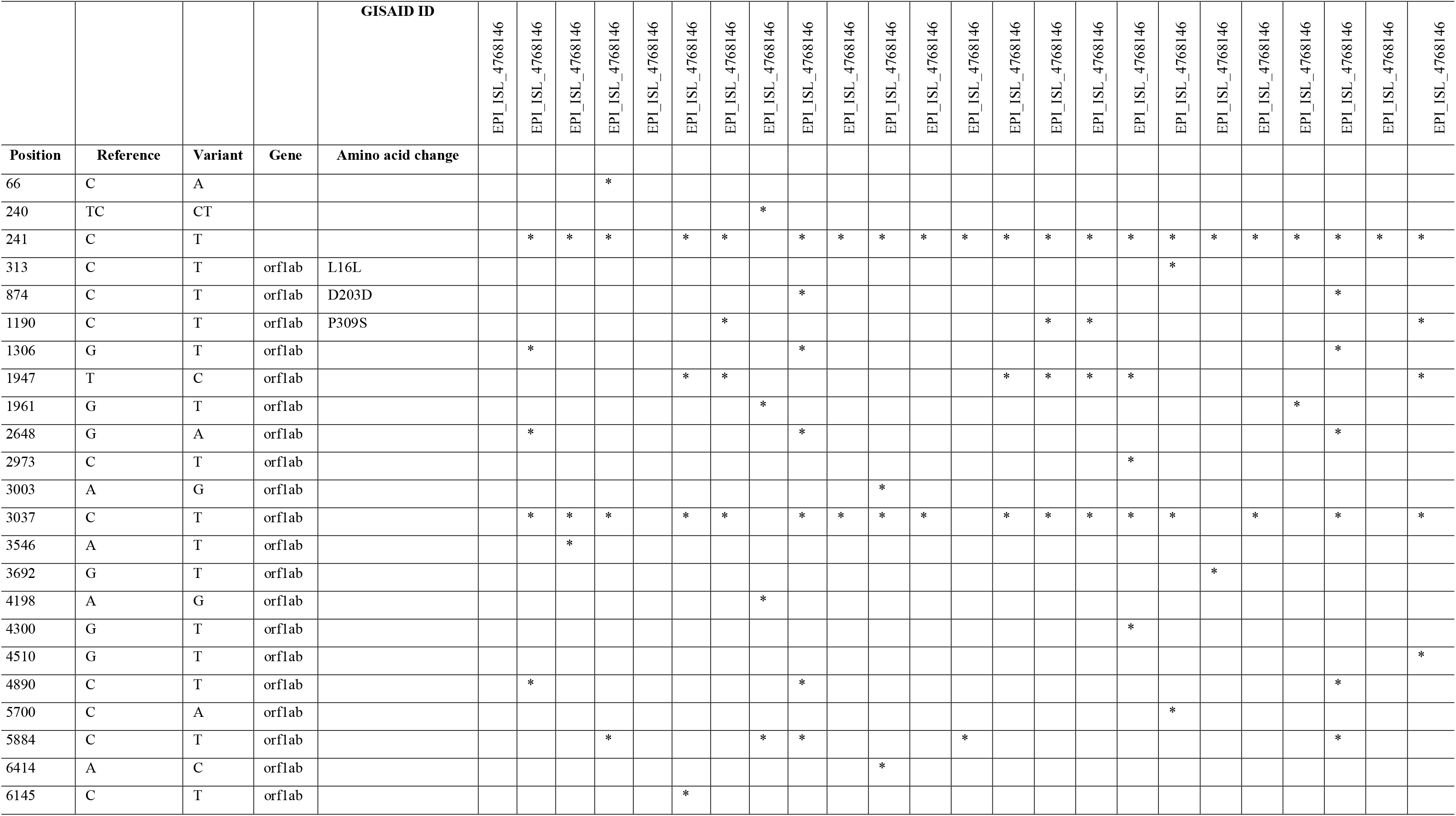
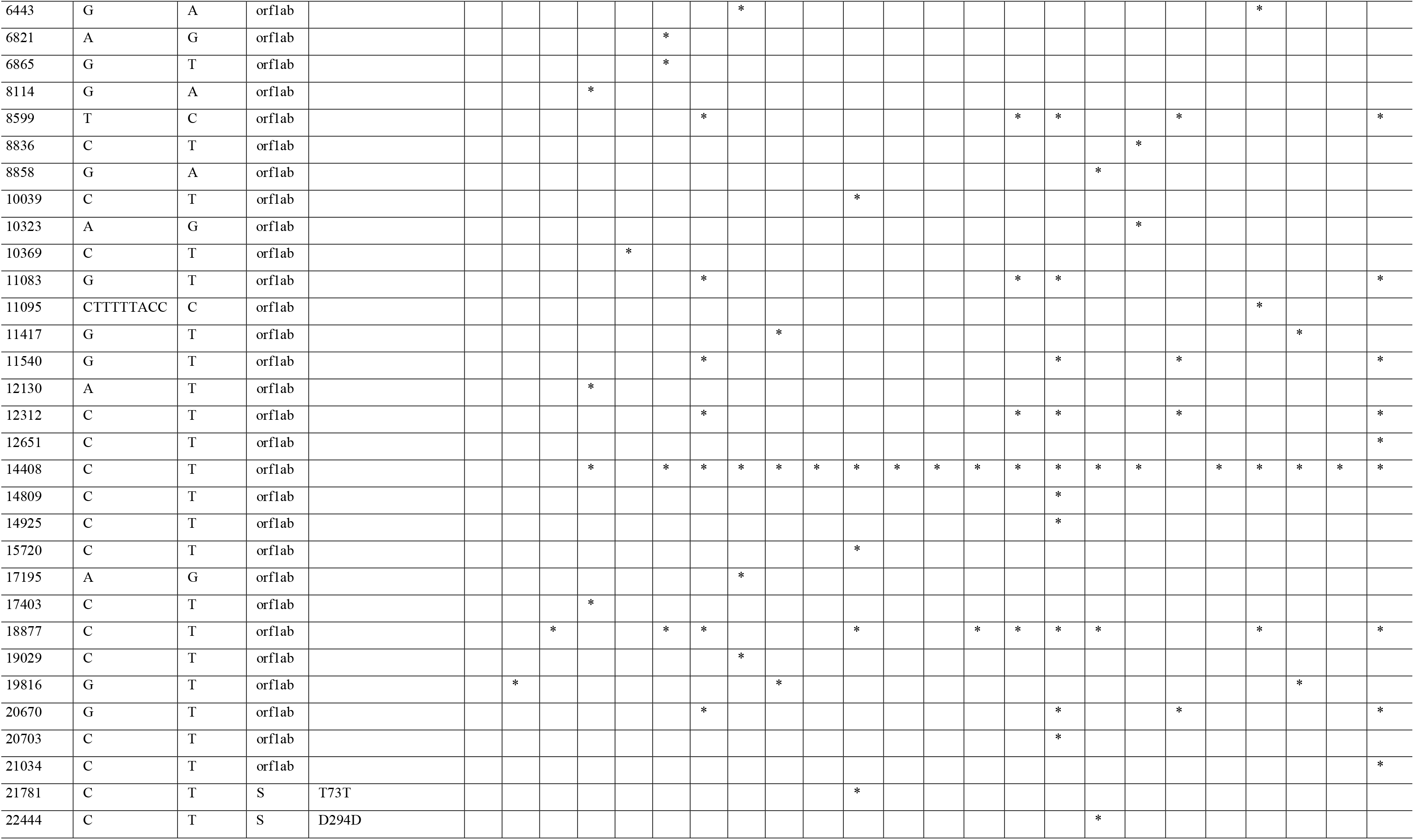
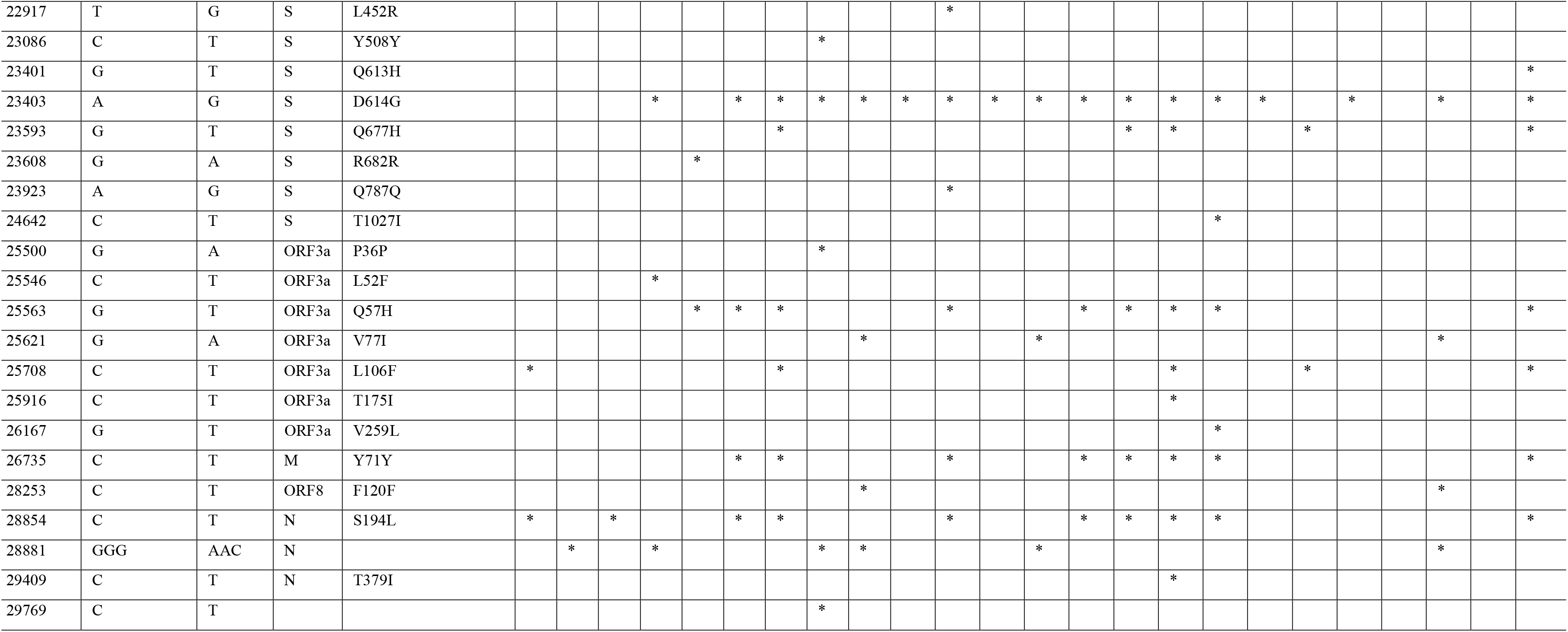
Mutations detected and their respective amino acid changes when compared to NCBI reference genome (SARS-CoV-2, Wuhan Hu-1, MN908947.3) (* marks indicate the presence of mutation)

Non-synonymous mutations were detected in ORF1ab, S, N, and ORF3a genes (Table 3). Mutations in the S gene such as D614G, L452R, Q613H, Q677H, T1027I have been identified as previously reported. Non-synonymous mutation D614G has been identified in most of the genome sequences, as reported widely in the literature that increases infectivity by adding more functional S protein into the virion causing more severity[21]. Other identified mutations are also linked with increase viral infectivity. Also several previously reported mutations like S194L in the N gene; Q57H, L106F, T175I in the ORF3a gene were observed. L52F, V77I mutations in ORF3a were also identified in one and three cases respectively.

A few possible novel mutations were also observed in ORF1ab, N, and ORF3a (Table 3). P309S mutation in the ORF1ab gene was identified in four samples. T379I mutation in the N gene was observed in one case. These mutations in the genome indicate the presence of multiple variations of the virus in Uttar Pradesh.

141 SARS-CoV-2 genome sequences from Lucknow, Uttar Pradesh were downloaded from GISAID to construct the phylogenetic tree with our 23 sequenced SARS-CoV-2 genome sequences. The sequenced 23 SARS-CoV-2 genome sequences were found in clade 20 A and 20B, out of which 16 variants were found related to clade 20 A, and the remaining 6 variants were found related to clade 20 B (Figure 1). Out of 23 genome sequences, 6 variants were identified as B.1 lineage, 8 variants as of B.1.36 lineage, 6 variants as of B.1.1.216 lineage and one variant was of B.1.456 lineage while the remaining two variants have not shown an association with any of lineages.

**Figure 1:**
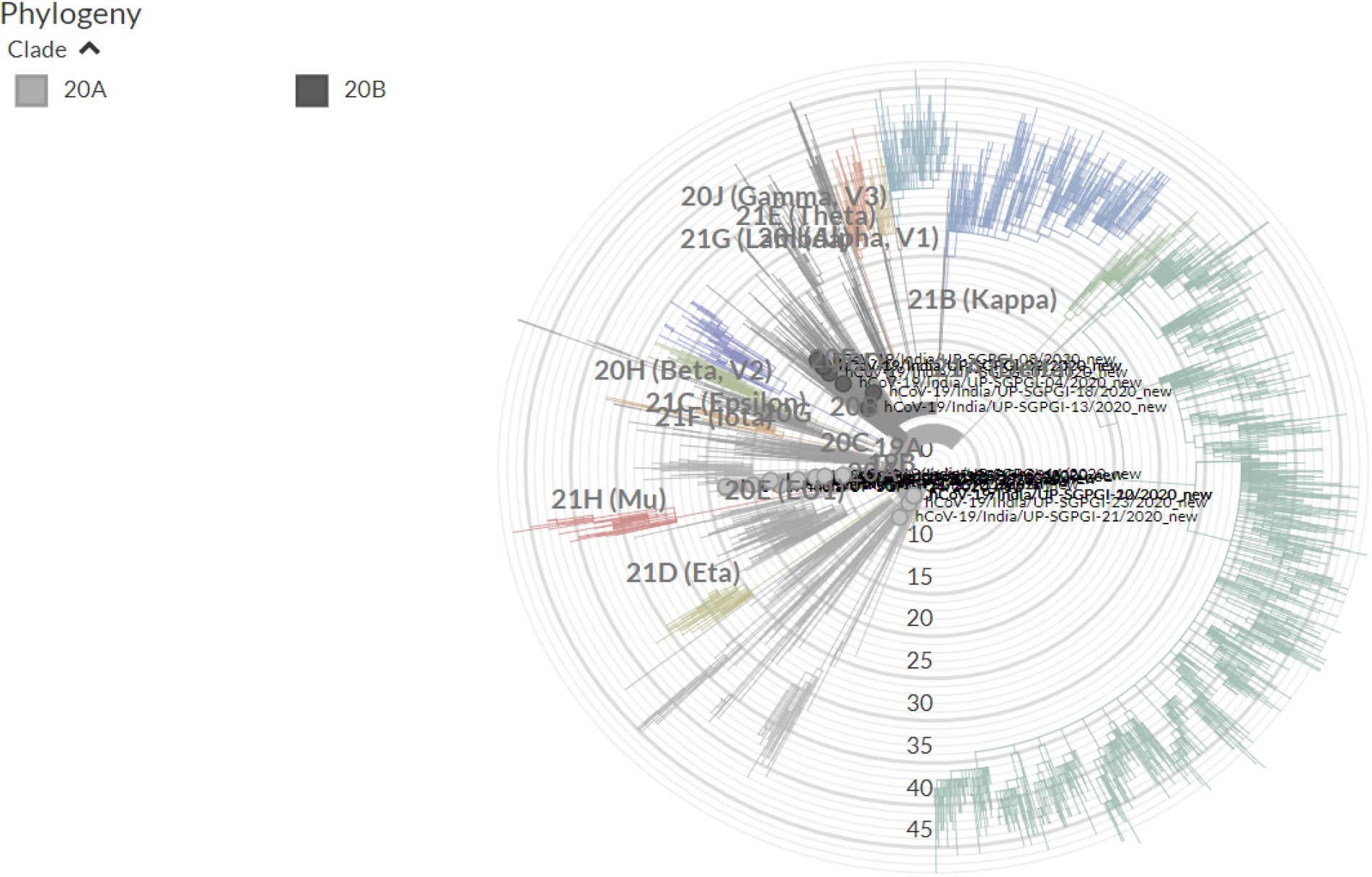
Phylogenetic analysis of 164 SARS-CoV-2 genome sequences: including 23 variants of present study and 141 variants from different regions of Lucknow, Uttar Pradesh, were retrieved from GISAID database. Nextclade was used for phylogentic analysis and nextstain nomenclature of all variants as shown in figure.

## Conclusion

We sequenced twenty-three SARS-CoV-2 genomes from the positive clinical samples collected during the first wave from Uttar Pradesh, India. Several identified mutations were already reported, while a few of the identified mutations could be novel. Most of the samples had D614G non-synonymous mutation. Phylogenetic analysis of the isolated viral genomes showed high similarities with the previously isolated SARS-CoV-2 genomes from Uttar Pradesh. Future studies can warranted to understand if these mutations potentially influence host susceptibility, pathogenicity, and virulence.

## Acknowledgment

The study was supported by intramural grants (A-24-PGI/IMP/81/2020) and overhead funds to ST. The authors wish to thank Dr. Suman Misra and Dr. Arvind Kumar (Department of Molecular Medicine, SGPGIMS, Lucknow) for their technical help.

